# Identification and characterization of long noncoding RNAs and their association with acquisition of blood meal in *Culex quinquefasciatus*

**DOI:** 10.1101/2020.03.06.980359

**Authors:** Azali Azlan, Mardani Abdul Halim, Faisal Mohamad, Ghows Azzam

## Abstract

The Southern house mosquito, *Culex quinquefasciatus* (*Cx. quinquefasciatus*) is an important vector that transmit multiple diseases including West Nile encephalitis, Japanese encephalitis, St. Louis encephalitis and lymphatic filariasis. Long noncoding RNAs (lncRNAs) involve in many biological processes such development, infection, and virus-host interaction. However, there is no systematic identification and characterization of lncRNAs in *Cx. quinquefasciatus*. Here, we report the first ever lncRNA identification in *Cx. quinquefasciatus*. By using 31 public RNA-seq datasets, a total of 4,763 novel lncRNA transcripts were identified, of which 3,591, 569, and 603 were intergenic, intronic, and antisense respectively. Examination of genomic features revealed that *Cx. quinquefasciatus* shared similar characteristics with other species such as short in length, low GC content, low sequence conservation, and low coding potential. Furthermore, compared to protein-coding genes, *Cx. quinquefasciatus* lncRNAs had lower expression values, and tended to be expressed in temporally-specific fashion. In addition, weighted correlation network and functional annotation analyses showed that lncRNAs may have roles in blood meal acquisition of adult female *Cx. quinquefasciatus* mosquitoes. This study presents the first systematic identification and analysis of *Cx. quinquefasciatus* lncRNAs and their association with blood feeding. Results generated from this study will facilitate future investigation on the function of *Cx. quinquefasciatus* lncRNAs.

## Introduction

The Southern house mosquito, *Culex quinquefasciatus* (*Cx. quinquefasciatus*), is an important member of the *Cx. pipiens* complex that spreads multiple types of diseases such as West Nile encephalitis, Japanese encephalitis, St. Louis encephalitis and lymphatic filariasis (Arensburger et al., 2010; Nchoutpouen et al., 2019). *Cx. pipiens* complex is composed of six members namely, *Cx. molestus* Forskll, *Cx. pipiens* Linneaus, *Cx. quinquefasciatus* Say, *Cx. pallens* Coquille, *Cx. globocoxitus* Dobrotworsky, and *Cx. australicus* Dobrotworsky & Drummond (Farajollahi et al., 2011; Zittra et al., 2016). *Cx. quinquefasciatus* mosquitoes are widespread and predominant in urban area because they are well-adapted to human made habitats (Nchoutpouen et al., 2019). Due to its important epidemiological role, *Cx. quinquefasciatus* has become a subject for many scientific studies. Several studies used next-generation sequencing to investigate genome-wide transcriptional response in *Cx. quinquefasciatus* upon blood meal acquisition (Reid et al., 2015; Taparia et al., 2017), insecticide exposure (Reid et al., 2012; Rezende et al., 2019), virus infections (Göertz et al., 2019; Paradkar et al., 2015; Rückert et al., 2019, 2019), and vector competence (Shin et al., 2014).

Many genes, especially protein-coding, have been implicated to be responsible in blood-feeding. The ability of *Cx. quinquefasciatus* to carry many harmful diseases is due to its blood-feeding behavior. Newly eclosed female mosquitoes need a minimum of 48 hours before they can feed on blood, and subsequently mate (Reid et al., 2015). For example, cytochrome P450s, proteases, and odorant-binding proteins were upregulated during blood-feeding period (Reid et al., 2015). Another study performed RNA-seq on antennae of blood-fed and nonblood-fed females, and found that chemosensory genes, odorant-binding proteins, ionotropic and odorant receptors were among the genes that were differentially expressed (Taparia et al., 2017). Besides blood meal, RNA-seq was also used to identify genes that may be involved upon insecticide exposure and resistance. Taken together, the utilization of RNA-seq approach enables the researcher to observe the differences in *Cx. quinquefasciatus* gene expression level at a genomic scale under different physiological conditions.

Even though protein-coding genes have always been the focus in transcriptomic studies, several reports showed that noncoding RNAs (ncRNAs) are implicated to play major roles in mosquitoes (Azlan et al., 2019b; Etebari et al., 2017, 2016; Göertz et al., 2019; Gu et al., 2013; Miesen et al., 2016; Rückert et al., 2019). Regulatory ncRNAs include small RNAs and long noncoding RNA (lncRNAs). Small RNA consists of micro-RNAs (miRNAs), short-interfering RNAs (siRNAs), and PIWI-interacting RNA (piRNAs), all of which regulate gene expression at the post-transcriptional level (Azlan et al., 2016; Brennecke et al., 2007; Coller and Parker, 2005; Djuranovic et al., 2012). lncRNAs, on the other hand, are RNA species of more than 200 nt in size that do not encode amino acids (Clark and Mattick, 2011; Ulitsky and Bartel, 2013). Due to next-generation sequencing and bioinformatics, identification of lncRNAs has been made possible in many organisms including mosquito such as *Aedes aegypti* (*Ae. aegypti*).

Previous study in *Ae. aegypti* showed that a subset of lncRNAs were highly expressed during early embryo (0-8 hour), and they were also enriched in blood-fed ovary. This suggests that lncRNAs may be maternally inherited, and they may play critical roles in early embryonic development (Azlan et al., 2019b). Beside embryonic development, lncRNAs in mosquitoes are also involved in host-virus interaction. For example, knockdown of *Ae. aegypti* lncRNA (lincRNA_1317) resulted in an increase in virus replication (Etebari et al., 2016). Although these studies suggest the regulatory roles of lncRNAs, the specific mechanism of lncRNAs functions are still unknown. Up until now, there has been no lncRNA studies in *Cx. quinquefasciatus*. We believed that, similar to *Ae. aegypti*, lncRNAs may possess certain regulatory roles in *Cx. quinquefasciatus*. In order to dissect their functional roles, it is important to first systematically annotate lncRNAs in the genome of *Cx. quinquefasciatus*.

Here, we report the first annotation of lncRNAs in *Cx. quinquefasciatus* genome using 31 publicly available RNA-seq data. We applied stringent bioinformatics analysis to confidently predict a total of 4,763 novel lncRNAs that correspond to 4,037 loci in the genome. We then characterized the newly identified lncRNAs such as expression dynamics, genomic features, and sequence conservation. We also investigate the transcriptional response of *Cx. quinquefasciatus* lncRNAs upon blood meal acquisition. Furthermore, through our co-expression network analysis, a subset of *Cx. quinquefasciatus* lncRNAs may participate in the taking and processing of blood meal.

## Materials and Methods

### Preparation of public RNA-seq datasets

31 RNA-seq datasets were downloaded from NCBI SRA websites. List of accession number can be found in **Supplementary Data 1**. FASTQC was used to check the quality of raw reads. Trimmomatic version 0.38 (Bolger et al., 2014) was used to clip adapters and low quality reads (<20 phred score).

### Alignment of RNA-seq reads to reference genome and transcriptome assembly

Cleaned reads were individually aligned to *Cx. quinquefasciatus* genome (CpipJ2, VectorBase) using Hisat2 (Kim et al., 2015). The reads were mapped according to the strandedness of the libraries. BAM files generated by Hisat2 were used as input for transcriptome assembly using Stringtie version 1.0.1 (Pertea et al., 2015). Annotation file of *Cx. quinquefasciatus* in GTF format (VectorBase) was used to guide the assembly. We set the minimum size of transcripts to be assembled was 200 bp. All Stringtie output files were merged into a single unified transcriptome assembly using Stringtie merge option. The resulting unified transcriptome assembly (GTF format) was compared to the reference annotation using Gffcompare (https://github.com/gpertea/gffcompare), and transcripts with class code “u”, “i”, and “x” were kept for lncRNA prediction analysis.

### lncRNA prediction

Transcripts with class code “u”, “i”, and “x” were subjected to TransDecoder analysis to predict long ORF, which was set to be minimum of 100 amino acids. Transcripts that do not have long ORF were then analyzed for their coding potential using CPC2 (Kang et al., 2017), PLEK (Li et al., 2014), and CNCI (Sun et al., 2013). Only transcripts that were categorized as noncoding by all three softwares were retained for downstream analyses. We then used BLASTX to find sequence similarity of the potential lncRNA transcripts against Swissprot database. Transcripts having high sequence similarity (E-value < 1e-3) were discarded. We then removed transcripts that had no strand information and located within less than 2 kb scaffold-end range.

### Transcript quantification and expression

Salmon version 0.10.1 (Patro et al., 2017) was used with default parameters to compute the expression of each transcript. We combined protein-coding and lncRNA transcripts together into a single FASTA file when running Salmon. TPM value from Salmon was used for downstream analysis.

### Validation of lncRNAs by reverse-transcribed PCR

About 20 adult females of VCRU-lab strain *Cx. quinquefasciatus* were obtained from Vector Control Research Unit, Universiti Sains Malaysia. The mosquitoes were homogenized for RNA isolation and subsequent reverse transcription for cDNA synthesis. For RNA isolation, Rneasy Mini Kit (Qiagen, cat. no: 74104) was used according to manufacturer’s protocol. Reverse transcription was carried out using iScript Reverse Transcription Supermix (Bio-Rad Laboratories, California; cat. no. 1708840) according to manufacturer’s protocol. We then performed PCR using the cDNA as templates. We designed PCR primers that span exon-exon junction using Primer Blast (https://www.ncbi.nlm.nih.gov/tools/primer-blast/). Primers used in this study is listed in **Supplementary Data 2**.

### Expression specificity

We used JS divergence method to compute expression specificity of lncRNAs and protein-coding genes. MATLAB version R2018b was used to calculate JS score using the formula previously described (Cabili et al., 2011).

### Co-expression analysis

WGCNA version 1.4.6 was used to perform co-expression analysis between lncRNAs and protein-coding genes (Langfelder and Horvath, 2008; Zhang and Horvath, 2005). We computed variance of each lncRNA and protein-coding gene, and ranked them in descending order. We only picked top 5000 genes for co-expression analysis.

### Functional annotation

g;Profiler was used for functional annotation and enrichment analysis (Reimand et al., 2007). g:SCS threshold was used for multiple testing correction.

## Result

### Identification of *Cx. quinquefasciatus* lncRNAs

In this study, we used 31 public RNA-seq datasets which were composed of approximately 849 million reads as raw inputs for lncRNA prediction. Public RNA-seq datasets used in the study were listed in **Supplementary Data 1**. Pipeline for lncRNA prediction can be found in **Figure 1**. Each library (paired or single-end) was individually mapped against *Cx. quinquefasciatus* genome (CpipJ2, VectorBase) using splice-aware aligner tool, Hisat2 (Kim et al., 2015). Each alignment file was then used for transcriptome assembly using Stringtie, and all 31 Stringtie output files were merged using Stringtie merge option (Pertea et al., 2015). We then compared our merged transcriptome assembly with reference annotation (CpipJ2.4, VectorBase) using gffcompare (https://github.com/gpertea/gffcompare). A total of 64,689 transcripts that correspond to 37,532 loci were assembled by Stringtie. Out of 64,689, 45,238 of them were multi-exon transcripts. We managed to recover all *Cx. quinquefasciatus* known transcripts (19,859) using Stringtie; hence, validating our transcriptome assembly.

**Figure 1:**
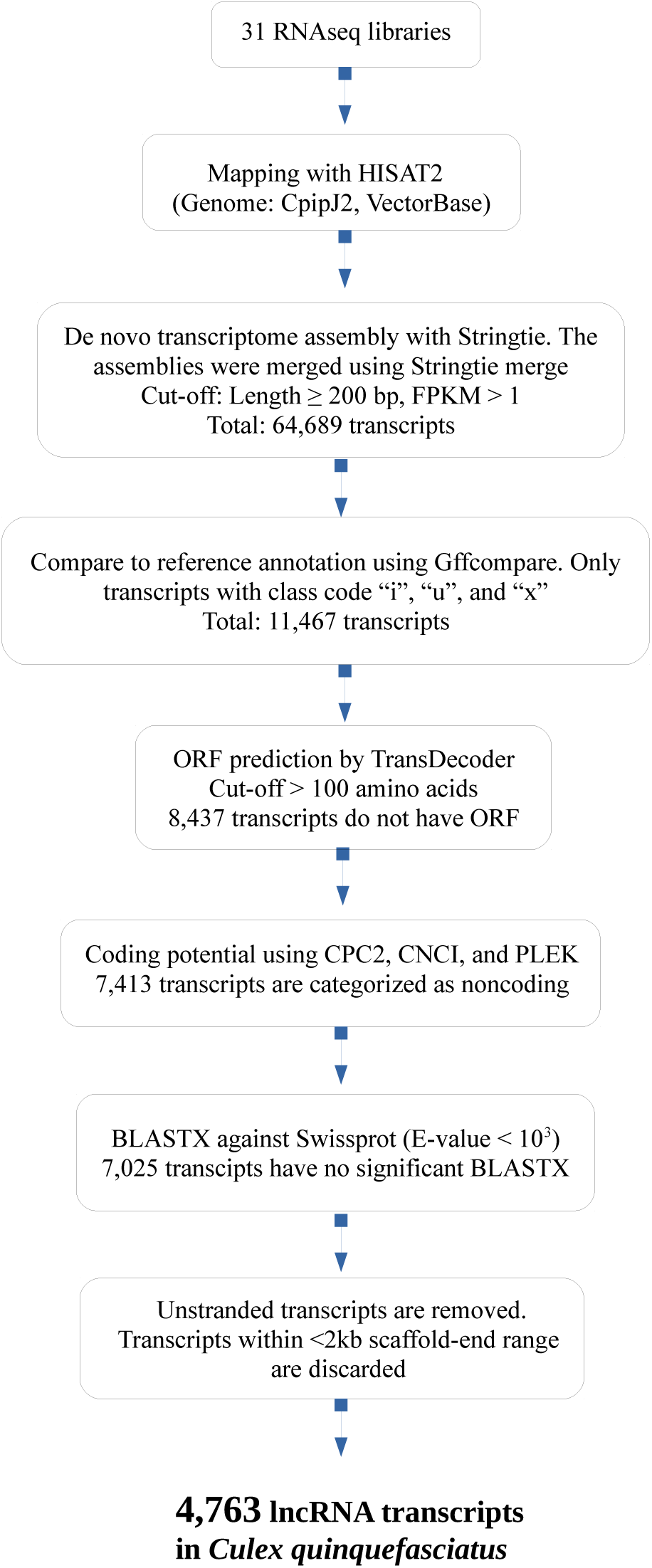
lncRNA identification. Summary of lncRNA prediction pipeline

After transcriptome assembly, we then performed stringent transcript filtering steps (**Figure 1**). Transcripts of less than 200 bp were discarded. We only chose transcripts with class code “i”, “u”, and “x”, all of which were given by gffcompare. Class code “i”, “u”, and “x” denote intronic, intergenic, and antisense to reference annotation respectively (https://github.com/gpertea/gffcompare). We discovered that a total of 11,467 transcripts were denoted as either “i”, “u” or “x”, and all of them were at least 200 bp in length. We then used TransDecoder (Haas et al., 2013) to predict long open-reading frame (ORF) within these transcripts. Out of 11,467, 2,994 transcripts were discarded because they were predicted to have ORF of more than 100 amino acids.

The remaining 8,473 transcripts were then evaluated for coding potential using CPC2 (Kang et al., 2017), PLEK (Li et al., 2014), and CNCI (Sun et al., 2013), and those classified as “noncoding” by all three algorithms were retained for downstream analysis. A total of 7,413 transcripts were found to be predicted as noncoding by all three softwares. To avoid false positive prediction, we used BLASTX (E-value cut-off 10-3) to find sequence similarity of those 7,413 transcripts with known proteins in Swissprot. The number of transcripts that have no significant BLASTX are 7,025. Finally, we removed unstranded transcripts, and transcripts located within less than 2kb scaffold-end range on the same strand. By applying this pipeline, we identified a total of 4,763 novel lncRNA transcripts, which derived from 4,037 distinct loci in the genome. From these 4,763 lncRNA transcripts, 3,591 of them were intergenic, while the remaining 569 and 603 transcripts were intronic and antisense to reference gene respectively. List of lncRNAs *Cx. quinquefasciatus* and their corresponding genomic loci can be found in **Supplementary Data 3**. We randomly selected 5 novel lncRNAs, and validated them through reverse-transcribed PCR (RT-PCR) using specific primers that span exon-exon junction (**Supplementary Figure 1**).

### Structural features of *Cx. quinquefasciatus* lncRNAs

Studies in other species showed that, when compared with protein-coding genes, lncRNAs displayed certain characteristics: shorter in size, lower GC content, low sequence conservation, and lower coding potential (Akay et al., 2019; Azlan et al., 2019b, 2019a; Etebari et al., 2016; Jenkins et al., 2015; Liu et al., 2017; Nam and Bartel, 2012; Wu et al., 2016; Young et al., 2012). We asked if this is also true for *Cx. quinquefasciatus* lncRNAs. We discovered that, in general, *Cx. quinquefasciatus* lncRNAs were significantly shorter than protein-coding transcript (Kolmogorov-Smirnov test (KS-test) *p*-value < 2.2e-16, **Figure 2a**). Mean of lncRNAs and protein-coding transcripts were 699 bp and 1,377 bp respectively. Meanwhile, lncRNAs had a median size of 463 whereas the median length protein-coding transcripts were 1,047 bp.

**Figure 2:**
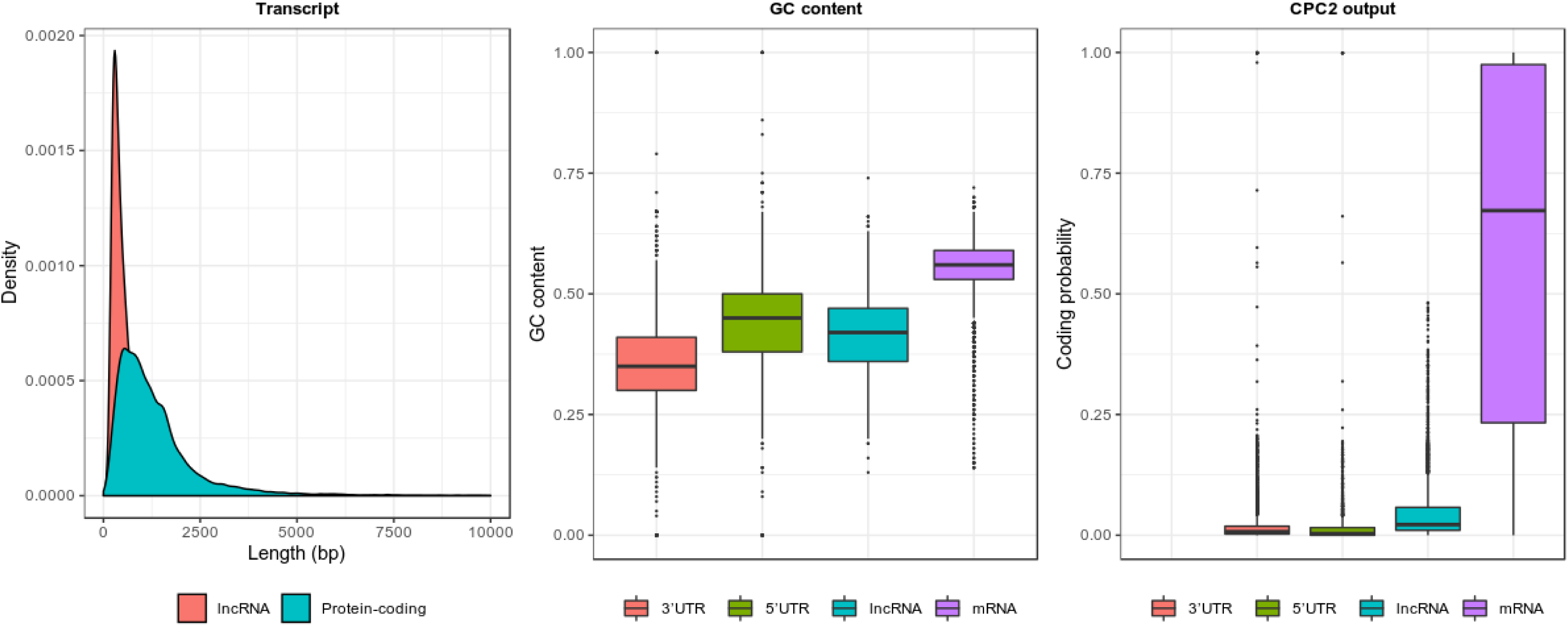
Features of *Cx. quinquefasciatus* lncRNAs. a) Distribution of transcript length b) GC content c) Coding potential from CPC2 algorithm

We then checked the GC content of lncRNAs and compared them with protein-coding transcripts, mRNAs. We also included other noncoding sequences sequences such as 3’UTR and 5’UTR. We found that, in general, noncoding sequences (lncRNAs, 3’UTR, and 5’UTR) have slightly lower GC content than mRNA (**Figure 2b**). Median GC content of lncRNA, 3’UTR, 5’UTR, and mRNA were 42%, 35%, 45%, and 56% respectively. In addition, mean GC content of noncoding sequences were lower than mRNA (mRNA: 55%, lncRNA: 41%, 5’UTR: 44%, 3’UTR: 36%). Beside GC content, we also examined coding potential of noncoding sequences and compared them with mRNA transcripts. CPC2 software showed that mRNA had the highest median and mean coding probability score (mean: 0.65 and median: 0.672). Median score for lncRNA, 5’UTR, and 3’UTR were 0.02, 0.004, and 0.007 respectively. Meanwhile, average probability of lncRNA, 5’UTR and 3’UTR were 0.047, 0.023, and 0.027 respectively.

Previous studies reported that lncRNAs have low sequence conservation (Azlan et al., 2019b; Etebari et al., 2016; Wu et al., 2016). Here, we assessed sequence conservation of the newly predicted *Cx. quinquefasciatus* lncRNAs using BLASTN against other insect genomes such as *Ae. aegypti, Aedes albopictus* (*Ae. albopictus*), *Anopheles gambiae* (*An. gambiae*), and *Drosophila melanogaster* (*D. melanogaster*). To determine the level of nucleotide similarity, we used bit score from BLASTN algorithm (Azlan et al., 2019b; Etebari et al., 2016). Similar to previous reports (Azlan et al., 2019b; Etebari et al., 2016; Wu et al., 2016), *Cx. quinquefasciatus* lncRNAs, in general, had lower sequence conservation than protein-coding transcripts (**Figure 3a**).

**Figure 3:**
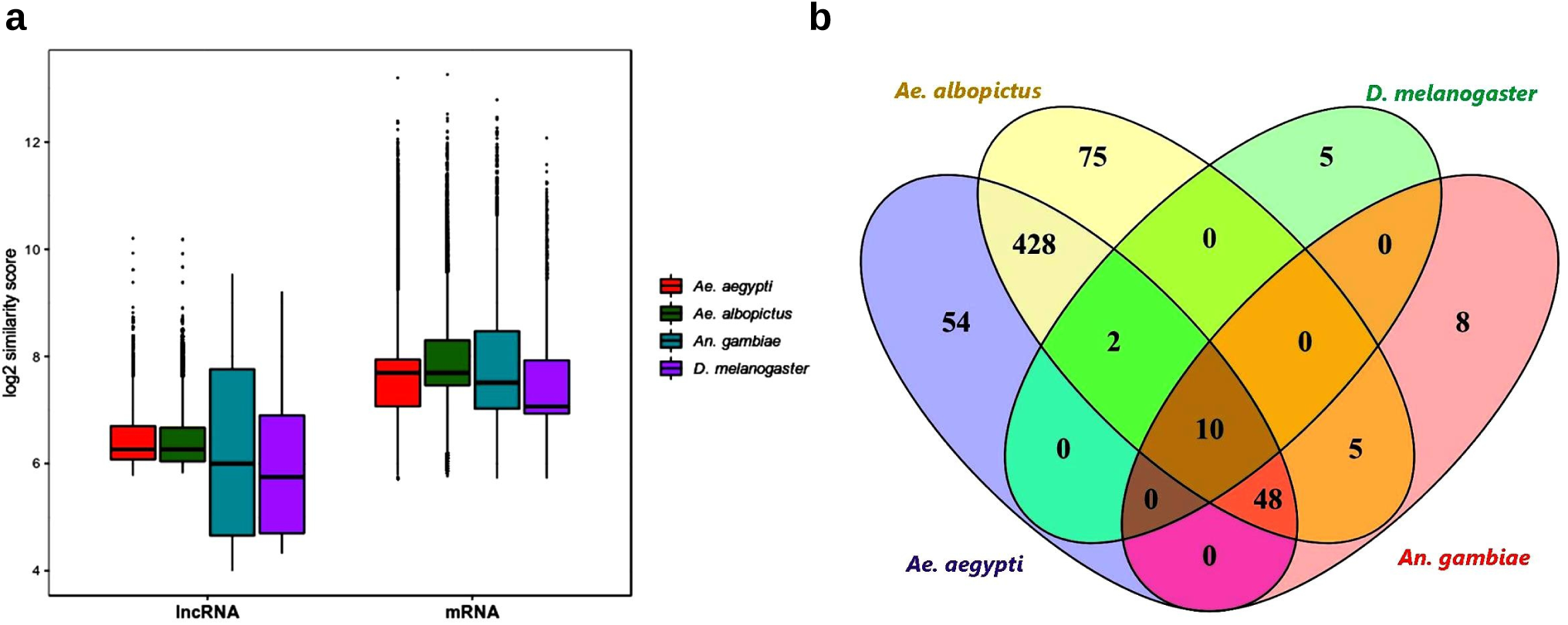
Sequence conservation of *Cx. quinquefasciatus* lncRNAs. a) Boxplot of similarity bit score using BLASTN algorithm. b) The Venn diagram shows the number of *Cx. quinquefasciatus* with E-value below 1e-5 in other species using BLASTN algorithm.

### *Cx. quinquefasciatus* lncRNAs are lowly expressed and temporally-specific

To assess the dynamics of lncRNA expression across different time points, we analyzed transcriptome data by Reid et al. (2015), which generated RNA-seq libraries of female mosquitoes *Cx. quinquefasciatus* at 2, 12, 24, 36, 48, 60 and 72 hour post-eclosion (Reid et al., 2015). We reanalyzed this time-course transcriptome data to compute the expression level of *Cx. quinquefasciatus* lncRNAs. We discovered that, across all time points, the expression level of lncRNA was lower than protein-coding genes (KS test *p*-value < 2.2e-16, **Figure 4a**).

**Figure 4:**
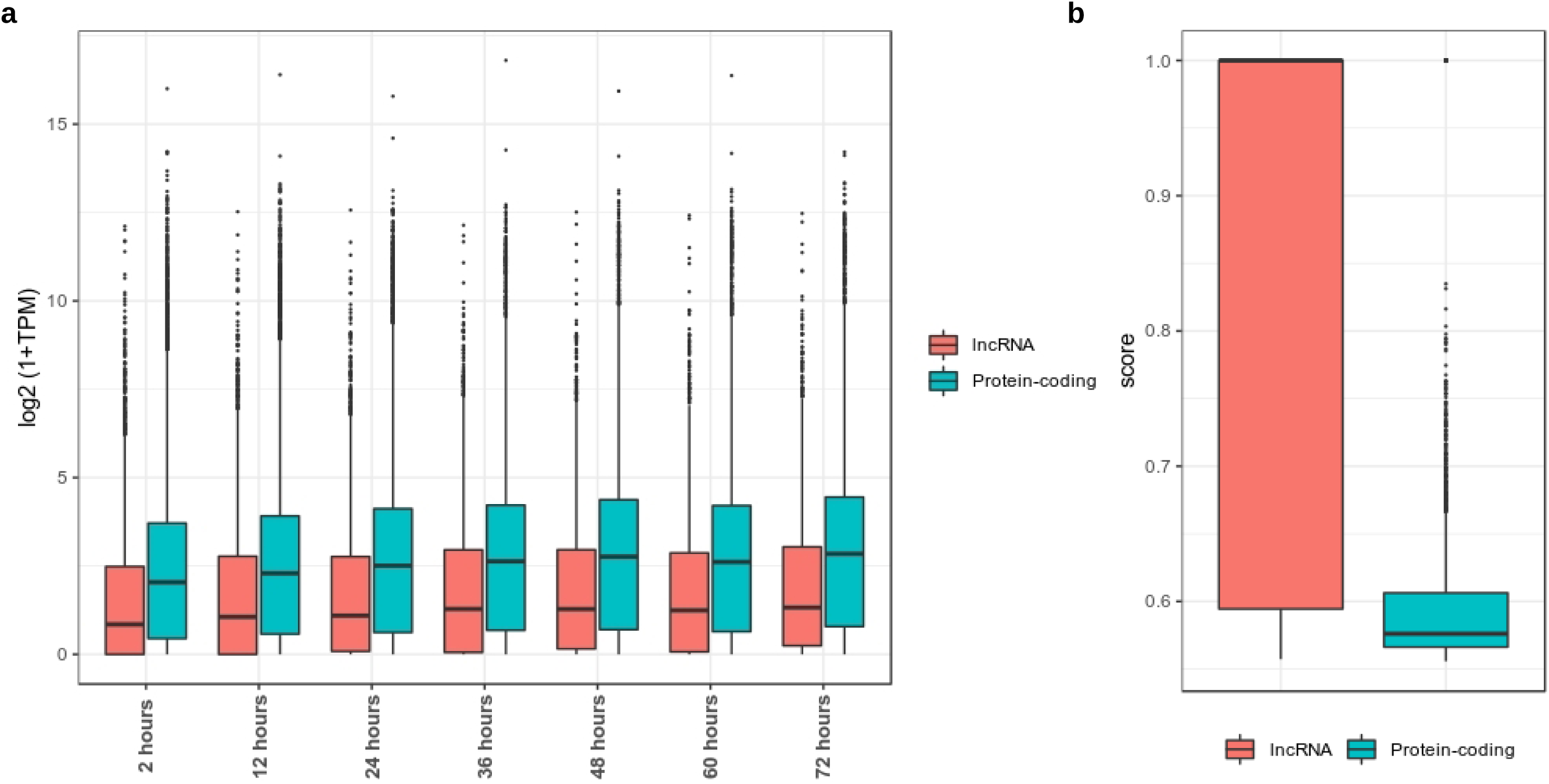
Expression of *Cx. quinquefasciatus* lncRNAs. a) Distribution of expression of lncRNAs and protein-coding genes across different time points. b) Distribution of specificity score of lncRNAs and protein-coding genes.

Previous studies reported that lncRNA in various organisms displayed a more temporally and tissue-specific pattern in expression (Azlan et al., 2019a, 2019b; Cabili et al., 2011; Wu et al., 2016). We then checked if *Cx. quinquefasciatus* lncRNAs have the same characteristics. To investigate expression specificity of lncRNAs, we employed Jensen-Shannon (JS) score, that ranges between zero to one (Cabili et al., 2011). This specificity score quantifies the similarity in expression value across different time points. Genes having JS score of one indicate the extreme case in which it is only expressed in one time point, whereas those having a score close to zero are ubiquitously expressed in all time points with relatively similar value of expression (Cabili et al., 2011). Similar to previous reports, the expression of *Cx. quinquefasciatus* lncRNAs was more temporally specific than protein-coding genes (KS test *p*-value < 2.2e-16, **Figure 4b**). We found that 58% of lncRNA transcripts had perfect specificity score of one, whereas only 15% of protein-coding transcripts had JS score of one.

### *Cx. quinquefasciatus* lncRNAs have potential roles in blood meal acquisition

To investigate the possible involvement of lncRNA in blood meal acquisition, we analyzed transcriptome data by Reid et al. (2015), which aimed to identify genes, especially protein-coding genes, that might be crucial for taking and processing the blood meal in adult female *Cx. quinquefasciatus* (Reid et al., 2015). They generated paired-end RNA-seq libraries of seven different time points after eclosion (2, 12, 24, 36, 48, 60, and 72 hours) of adult females in order to examine temporal gene expression of post eclosion and prior to blood-feeding (Reid et al., 2015). They showed that adult females *Cx. quinquefasciatus* need at least 48 hour post-eclosion before they can take their first blood meal (Reid et al., 2015). Therefore, prior to 48 hours, it was hypothesized that genes necessary for blood meal acquisition would be differentially expressed, and after 48 hours, the differentially expressed genes would be important for processing the blood meal. We hypothesized that, besides protein-coding genes, lncRNAs in *Cx. quinquefasciatus* would also play regulatory roles in blood-feeding process.

Weighted Gene Correlation Network Analysis (WGCNA) has always been used to predict potential roles of lncRNAs in many species (Azlan et al., 2019a; Langfelder and Horvath, 2008; Wu et al., 2016; Zhang and Horvath, 2005). WGCNA can be used to search for clusters or modules of highly correlated genes, which in this case, correlation between lncRNAs and protein-coding genes (Langfelder and Horvath, 2008; Zhang and Horvath, 2005). To predict the potential roles of lncRNAs in blood meal acquisition, we used RNA-seq data from Reid et al. 2015, and performed WGCNA analysis on lncRNAs and protein-coding genes (Reid et al., 2015). WGCNA analysis yielded a total of 21 modules, and 7 of them exhibited strong and statistically significant correlation (correlation > 0.8, *p*-value < 0.05) with specific time points **(Figure 5)**. Module size ranged from 22 to 611, and the number of protein-coding genes and lncRNA in each module can be found in Table 1.

**Table 1:** Number of lncRNA and protein-coding genes in each module (Correlation > 0.8, P-value < 0.05)

| Modules | lncRNA | Protein-coding | Hour post-eclosion | Correlation (P-value) |
| --- | --- | --- | --- | --- |
| Green | 66 | 226 | 2 hours | 0.81 (0.03) |
| Brown | 103 | 508 | 2 hours | 0.99 (2e-5) |
| Magenta | 43 | 59 | 12 hours | 0.97 (2e-4) |
| Black | 41 | 150 | 24 hours | 0.84 (0.02) |
| Royal blue | 17 | 7 | 36 hours | 0.98 (6e-5) |
| Light yellow | 16 | 13 | 48 hours | 0.99 (1e-5) |
| Dark red | 12 | 10 | 60 hours | 0.99 (4e-5) |

**Figure 5:**
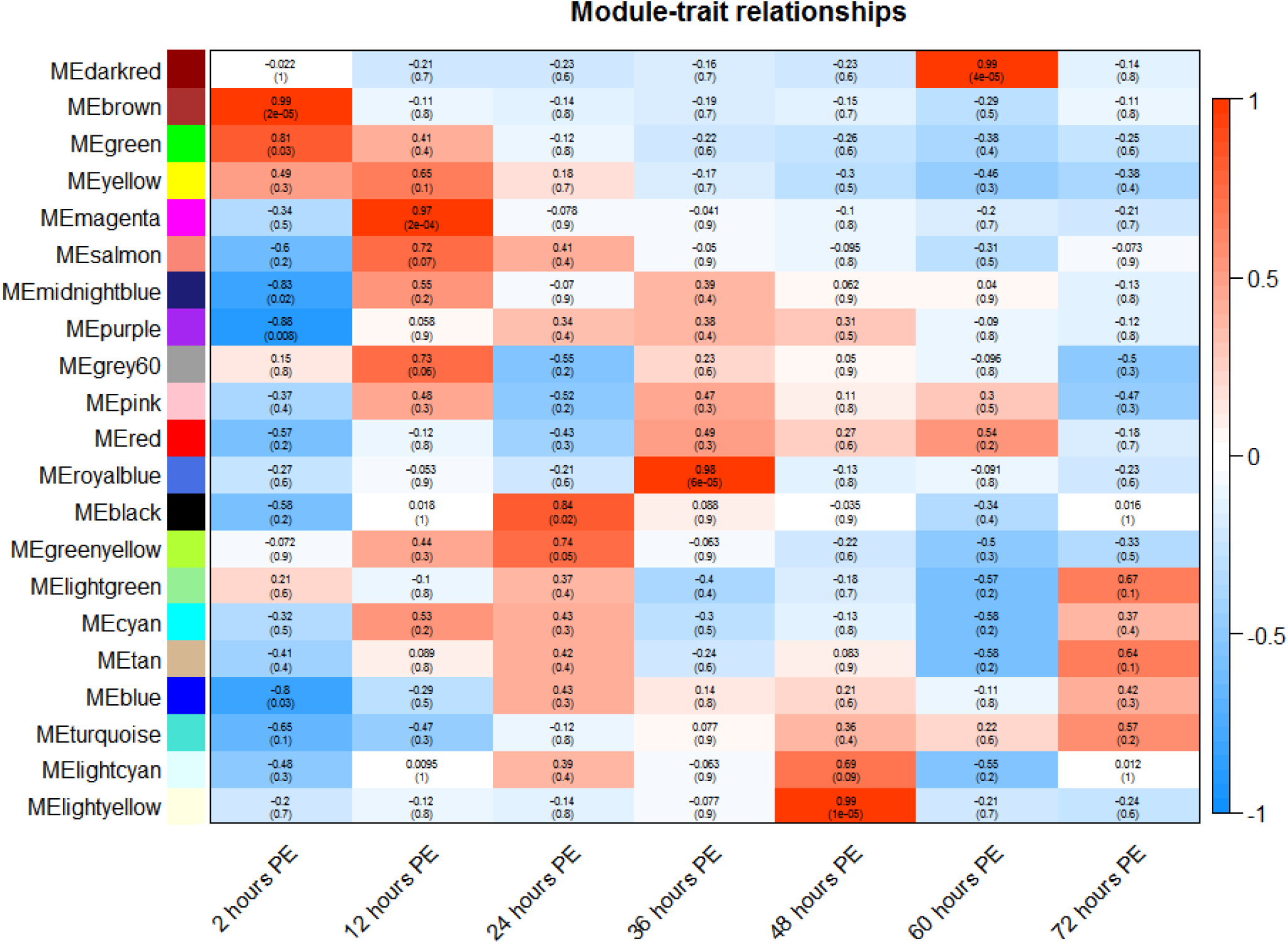
Module-trait correlation. Module-trait correlation and corresponding p-values was computed using WGCNA package. Red indicated positive correlation (correlation > 0) whereas blue color indicated negative correlation (correlation < 0). PE: post-eclosion.

Two modules were found to be significantly associated with 2 hours post-eclosion, and both of them had the among the largest number of genes within the clusters **(Table 1)**. Functional annotation using g:Profiler of these two modules revealed that lncRNAs at 2 hours had potential roles in diverse processes such as compound metabolic pathways, biosynthesis, catabolic process, and proteolysis (**Supplemental Data 4**). At 12 hour post-eclosion, several genes such as salivary proteins and cytochromes were found to be differentially expressed (Reid et al., 2015). We next asked if the same set of genes with similar functions could be found in our magenta module that was highly associated with 12 hours post-eclosion (correlation: 0.97, *p-*value: 2e-4). We found that several protein-coding genes within magenta module were salivary proteins and cytochromes, such as proline-rich salivary peptide (CPIJ007453), threonine-rich salivary mucin (CPIJ010046), cytochrome c oxidase assembly factor (CPIJ010284) and cytochrome P450 (CPIJ011837). Since 43 lncRNAs were clustered together with 59 protein-coding genes within magenta module, we hypothesized that this set of lncRNAs may participate in the same function as protein-coding genes. Therefore, results from WGCNA corroborated with previous findings (Reid et al., 2015), and suggest that lncRNA may intimately function together with protein-coding genes in order to physiologically prepare adult females *Cx. quinquefasciatus* for blood feeding and blood meal processing.

## Discussion

In this study, we performed genome-wide identification and characterization of lncRNAs in *Cx. quinquefasciatus* genome using a total of 31 relatively high-depth publicly available RNA-seq libraries. We presented a set of 4,763 novel lncRNAs transcripts which derived from 4,037 loci in *Cx. quinquefasciatus* genome. RNA-seq libraries used here were generated from many tissues and developmental stages, making our prediction pipeline to be relatively robust and comprehensive. We applied stringent filtering steps in our pipeline to reduce false positive and false negative prediction of non-coding transcripts (Azlan et al., 2019a, 2019b; Etebari et al., 2016; Nam and Bartel, 2012; Wu et al., 2016). The filtering steps include using more than one coding potential assessment algorithms and removing transcripts with no strand information (**Figure 1**). Taken together, lncRNA prediction pipeline used in this study is comparably equivalent to previous studies in other organisms, and it is

Analysis of genomic features revealed that lncRNAs identified in our study showed known characteristics of lncRNAs from other species. The characteristics are shorter in size, low sequence conservation, and lower GC content. The majority of *Cx. quinquefasciatus* lncRNAs shared high degree of sequence similarity with *Ae. albopictus* and *Ae. aegypti*, followed by *An. gambiae* and *D. melanogaster*. This lncRNA sequence similarity pattern concur with previous phylogenetic relationship study on known mosquitoes which showed that *Ae. albopictus* and *Ae. aegypti* are more evolutionarily closer to *Cx. quinquefasciatus* than *An. gambiae* (Chu et al., 2018). A total of 10 *Cx. quinquefasciatus* lncRNAs shared high sequence similarity with all insects (**Figure 3b**), suggesting that they are evolutionarily conserved and may play vital roles in insect development or basic cellular functions.

Investigation on the expression across different time points post-eclosion revealed that, compared to protein-coding genes, *Cx. quinquefasciatus* lncRNAs were more temporally specific and have lower expression level. This high temporal specificity indicates that lncRNAs have a smaller time window of expression that protein-coding genes. Even though lncRNAs have lower expression that protein-coding genes, their temporal specificity in expression suggests that they putatively execute specific biological functions at specific time point in development.

Furthermore, co-expression analysis by WCGNA revealed that *Cx. quinquefasciatus* lncRNAs were significantly correlated with specific time-point of pre-blood feeding immediately after eclosion, suggesting that lncRNAs may possess potential roles in blood meal acquisition by adult female mosquitoes. This blood meal acquisition by adult female is physiologically important as it is not only required for reproduction and mating, but it serves as a gateway for pathogens, making *Cx. quinquefasciatus* one of the most harmful mosquito vectors on Earth. However, the data presented here is simply descriptive, and it does not provide empirical evidence via experimentation on the specific mechanisms of lncRNA functions in blood meal acquisition. We believed that results generated in this study can be a starting point for dissecting the mechanism of *Cx. quinquefasciatus* lncRNAs functions. In conclusion, this research provides the first comprehensive annotation and characterization of *Cx. quinquefasciatus* lncRNAs, and their putative roles in blood-feeding. We believed that data presented here will be a valuable resource for genetics and molecular studies of ncRNAs in *Cx. quinquefasciatus*.

## Acknowledgment

We would like to thank all our collaborators and colleagues for the discussion and the work conducted in this lab. We would especially like to thank the Vector Control Research Unit, Universiti Sains Malaysia for providing the mosquito samples. This study was funded by the Universiti Sains Malaysia Research University Grant (1001/PBIOLOGI/811320) and MESTECC ScienceFund (305/PBIOLOGI/613238).

## Figure Legend

**Supplementary Figure 1:**
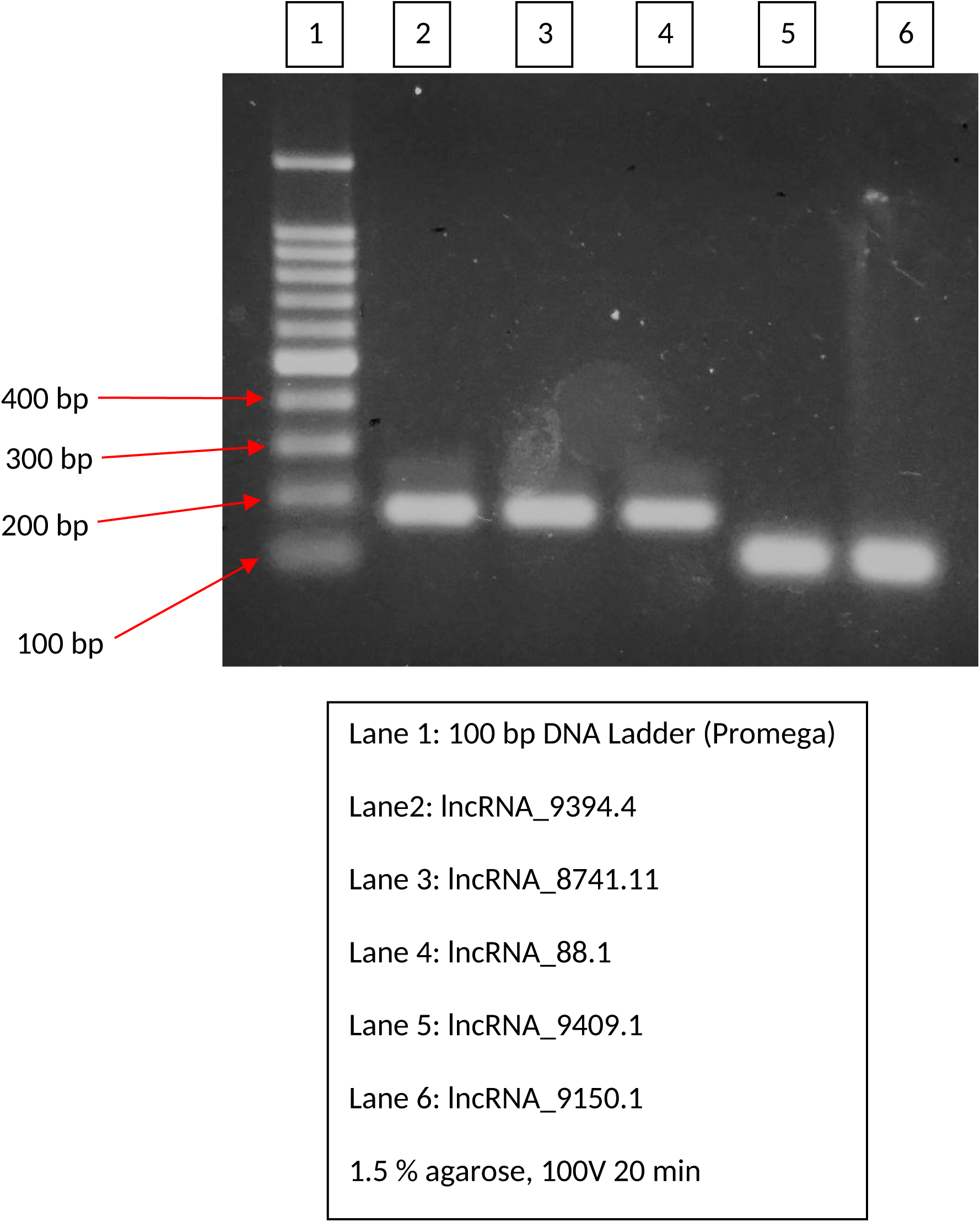
PCR products in 1.5% agarose gel.

**Supplementary Data 1:** List of 31 publicly available RNA-seq libraries used in this study

| Accession number | Condition | Single/Paired | Reference/Publication |
| --- | --- | --- | --- |
| SRR5500831 | sugar-fed replicate 2 | Single | Taparia et al., 2017 |
| SRR5500832 | sugar-fed replicate 1 | Single |  |
| SRR5500833 | blood-fed replicate 2 | Single |  |
| SRR5500834 | blood-fed replicate 1 | Single |  |
| SRR1005577 | adult female 2 hour post-eclosion | Paired | Reid et al., 2015 |
| SRR1005578 | adult female 12 hour post-eclosion | Paired |  |
| SRR1005579 | adult female 24 hour post-eclosion | Paired |  |
| SRR1005580 | adult female 36 hour post-eclosion | Paired |  |
| SRR1005581 | adult female 48 hour post-eclosion | Paired |  |
| SRR1005582 | adult female 60 hour post-eclosion | Paired |  |
| SRR1005583 | adult female 72 hour post-eclosion | Paired |  |
| SRR1539388 | control uninfected cell line | Paired | Paradkar et al., 2015 |
| SRR1539389 | control uninfected cell line | Paired |  |
| SRR1539390 | cell infected with West Nile virus | Paired |  |
| SRR1539391 | cell infected with West Nile virus | Paired |  |
| SRR9617919 | Susceptible colony | Paired | Rezende et al., 2019 |
| SRR9617921 | Susceptible colony | Paired |  |
| SRR9617922 | Susceptible colony | Paired |  |
| SRR9617920 | Resistant colony | Paired |  |
| SRR9617923 | Resistant colony | Paired |  |
| SRR9617924 | Resistant colony | Paired |  |
| SRR407541 | Egg | Single | - |
| SRR407542 | Third instar larvae | Single |  |
| SRR407543 | Pupae | Single |  |
| SRR407545 | Adult | Single |  |
| SRR407546 | S-lab strain third instar larvae | Single |  |
| SRR1050976 | Adult female | Paired | - |
| SRR1050983 | Adult female | Paired |  |
| SRR1462324 | Adult female | Paired | - |
| SRR991016 | Female antennae | Paired | - |
| SRR991017 | Female hind leg | Paired |  |

**Supplementary Data 2:**
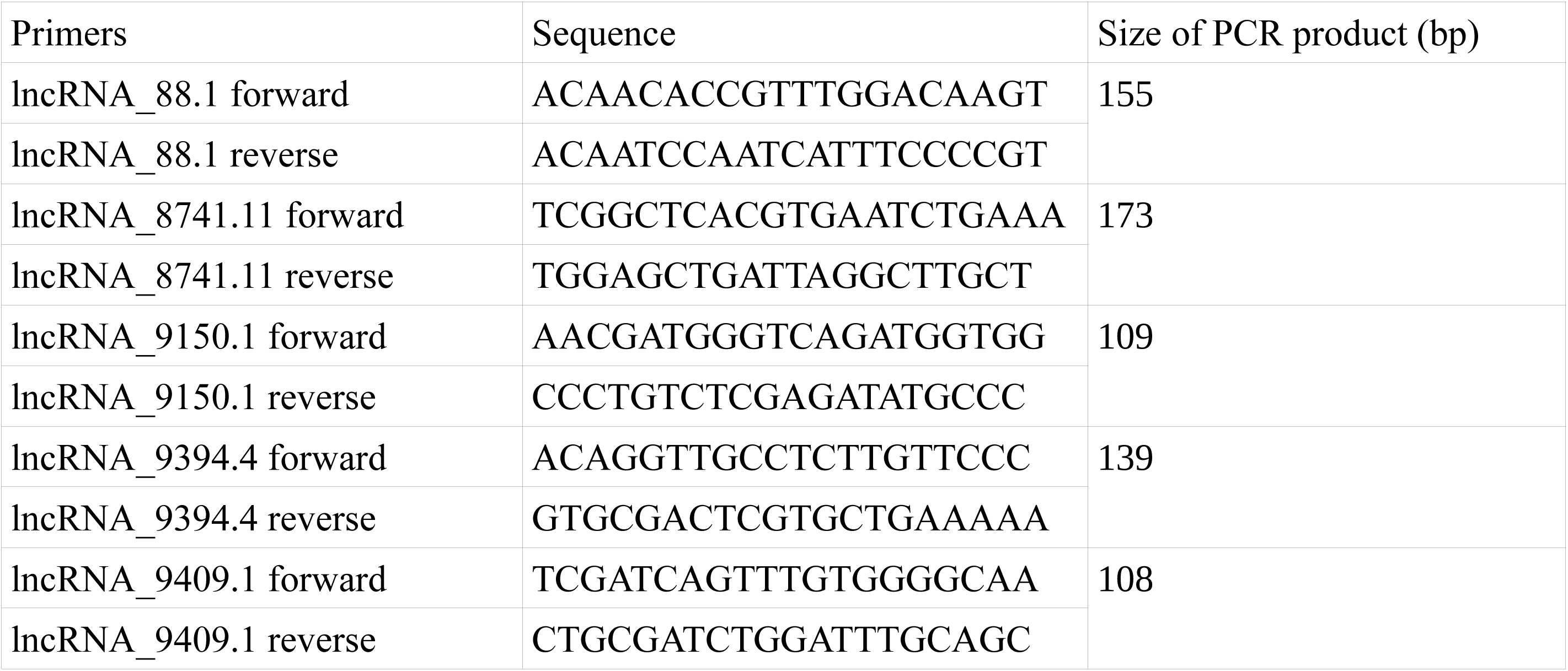
Primers used in this study

**Supplementary Data 3: List of lncRNAs in *Cx. quinquefasciatus* genome**

**Supplementary Data 4: Gene ontology and enrichment from g:Profiler**

